# Experimental support that natural selection has shaped the latitudinal distribution of mitochondrial haplotypes in Australian *Drosophila melanogaster*

**DOI:** 10.1101/103606

**Authors:** M. Florencia Camus, Jonci N. Wolff, Carla M. Sgrò, Damian K. Dowling

## Abstract

Cellular metabolism is regulated by enzyme complexes within the mitochondrion, the function of which are sensitive to the prevailing temperature. Such thermal sensitivity, coupled with the observation that population frequencies of mitochondrial haplotypes tend to associate with latitude, altitude or climatic regions across species distributions, led to the hypothesis that thermal selection has played a role in shaping standing variation in the mitochondrial DNA (mtDNA) sequence. This hypothesis, however, remains controversial, and requires evidence that the distribution of haplotypes observed in nature corresponds with the capacity of these haplotypes to confer differences in thermal tolerance. Specifically, haplotypes predominating in tropical climates are predicted to encode increased tolerance to heat stress, but decreased tolerance to cold stress. We present direct evidence for these predictions, using mtDNA haplotypes sampled from the Australian distribution of *Drosophila melanogaster*. We show that the ability of flies to tolerate extreme thermal challenges is affected by sequence variation across mtDNA haplotypes, and that the thermal performance associated with each haplotype corresponds with its latitudinal prevalence. The haplotype that predominates at low (subtropical) latitudes confers greater resilience to heat stress, but lower resilience to cold stress, than haplotypes predominating at higher (temperate) latitudes. We explore molecular mechanisms that might underlie these responses, presenting evidence that the effects are in part regulated by SNPs that do not change the protein sequence. Our findings suggest that standing variation in the mitochondrial genome can be shaped by thermal selection, and could therefore contribute to evolutionary adaptation under climatic stress.

## Introduction

The mitochondria are essential for eukaryote evolution, taking centre-stage in the process of cellular respiration. This process is regulated via a series of finely-coordinated interactions between the genes of two obligate genomes – nuclear and mitochondrial (Rand, et al. 2004; Wolff, et al. 2014). Indeed, because of the strong dependence of cellular respiration on mitochondrial-encoded gene products, biologists traditionally assumed that strong purifying selection would prevent any “function-encoding” genetic variation from accumulating within the mitochondrial DNA (mtDNA) (Ballard and Kreitman 1994; Rand 2001; Dowling, et al. 2008). The assumption of selective neutrality has, however, been challenged over the past decade via analyses of polymorphism and divergence data within the mtDNA sequences of metazoans. These analyses have used McDonald-Kreitman or similar tests of selection at the molecular level, to uncover signatures of recurrent adaptive evolution within the mitochondrial genome (Bazin, et al. 2006; James, et al. 2016). These have been complemented by studies using experimental approaches with the power to partition mitochondrial from nuclear genetic effects, which have demonstrated that the intra-specific genetic variation that exists within the mitochondrial genome commonly affects the expression of phenotypic traits, from morphological, to metabolic, to life-history (Rand 2001; Dowling, et al. 2008; Burton, et al. 2013; Dobler, et al. 2014).

Indeed, several lines of empirical evidence have emerged that support a novel hypothesis, which posits that the standing genetic variation that delineates the mtDNA haplotypes of spatially-disjunct populations has been shaped by natural selection imposed by the prevailing thermal climate (Mishmar, et al. 2003; Ballard and Whitlock 2004; Ruiz-Pesini, et al. 2004; Wallace 2007; Dowling 2014). The first support for this *mitochondrial climatic adaptation* hypothesis was provided by studies of mtDNA variation in humans, where patterns of amino acid variation were observed to align closely to particular climatic regions (Mishmar, et al. 2003; Ruiz-Pesini, et al. 2004), and where levels of genetic divergence between mtDNA haplotypes of different populations were shown to correlate with temperature differences between these populations (Balloux, et al. 2009). These studies on human mtDNA sequences are intriguing, but have in some cases proven difficult to replicate with different or larger datasets (Kivisild, et al. 2006; Sun, et al. 2007).

Additional support for the hypothesis has been provided from studies of other metazoans, some of which have reported signatures of positive selection on mtDNA genes sampled from populations inhabiting particular thermal environments (Foote, et al. 2011; Silva, et al. 2014; Ma, et al. 2015; Morales, et al. 2015), and others which have documented variation in mitochondrial gene or haplotype frequencies along clinal gradients associated with climate, such as latitude (Silva, et al. 2014; Consuegra, et al. 2015), altitude (Fontanillas, et al. 2005; Cheviron and Brumfield 2009), or with temperature itself (Quintela, et al. 2014). Such clinal patterns are, however, based on correlations between haplotype frequencies and environmental gradients. The possibility remains these correlations could be explained by neutral demographic processes, such as by multiple colonisations from different origins into different locations followed by admixture, sex-specific patterns of dispersal and introgression (given that the mtDNA sequence is maternally-inherited), or by recurrent occurrences of secondary contact (Endler 1977; Toews and Brelsford 2012; Adrion, et al. 2015; Bergland, et al. 2016).

Finally, support has been provided through laboratory experiments in invertebrates, which have reported that the expression of life-history phenotypes (Dowling, et al. 2007; Arnqvist, et al. 2010; Hoekstra, et al. 2013; Wolff, et al. 2016), as well as the transmission dynamics (Matsuura, et al. 1997; Doi, et al. 1999), associated with particular mtDNA haplotypes, or combinations of mitochondrial and nuclear (mito-nuclear) genotype, often depend on the thermal environment in which the study subjects are assayed. These studies have thus indicated that mitochondrial genetic variation is sensitive to thermal selection, at least when measured in the laboratory. However, these studies also have some caveats, given they are based on ‘between population’ (i.e, the mtDNA haplotypes used were drawn from distinct populations), or ‘between species’ (mtDNA from distinct species) experimental designs (Dowling, et al. 2007; Arnqvist, et al. 2010; Wolff, et al. 2016) (Matsuura, et al. 1997; Doi, et al. 1999; Hoekstra, et al. 2013). Inter-population and inter-species designs will presumably maximise the opportunity to detect mitochondrial genetic effects on organismal phenotypes, given that levels of mitochondrial divergence will increase with at inter-population and inter-species scales. Yet, the results obtained via such designs are less straightforward to interpret within the broader context of thermal adaptation, given that the most relevant level at which natural selection acts is on standing variation in mtDNA haplotypes within a population of a given species.

Accordingly, in this study we sought to directly test the *mitochondrial climatic adaptation* hypothesis, within the Australian distribution of the vinegar fly, *Drosophila melanogaster*. This species invaded Australia over a century ago (Hoffmann and Weeks 2007), and it is thought the Australian population was established from multiple introductions of flies from two origins; Africa and Eurasia (David and Capy 1988; Singh and Long 1992). A recent study of nuclear genome-wide allele frequencies from Australian populations concurs with this conclusion, with flies sampled from high latitudes closely related to cold-adapted European populations, and those from low latitudes more closely related to African populations (Bergland, et al. 2016). This study therefore provides a cautionary note, by indicating that colonisation history might well contribute to the existence of any latitudinal patterns in mtDNA haplotype frequencies that occur within Australia, rather than thermal selection acting on standing variation in mtDNA haplotypes (Adrion, et al. 2015; Bergland, et al. 2016).

Direct experimental evidence for the mitochondrial climatic adaptation hypothesis therefore requires a two-step approach. Firstly, evidence of shifts in the frequencies of mtDNA haplotypes along a gradient that aligns closely to the environment (e.g. latitude); and secondly, experimental evidence that links thermal sensitivities of these haplotypes when measured under controlled conditions in the lab, to their spatial distributions in the field. This has never previously been achieved for the genetic variation that resides within the mitochondrial genome. Indeed, when it comes to the evolutionary significance of clinal variation in general, there are surprisingly few examples in which latitudinal variation in allele frequencies has been linked clearly to variation in fitness (Adrion, et al. 2015).

## Results and Discussion

We collected field-inseminated female flies from 11 populations along an eastern Australian latitudinal cline (Table S1), and used these flies to initiate isofemale lines (lines initiated by a solitary gravid female), and ultimately mass-bred populations per latitudinal location (with each population kept in independent duplicate). Previous research has shown linear associations between the expression of thermal tolerance phenotypes, and allele frequencies of underlying candidate nuclear genes, along this latitudinal cline (Hoffmann, et al. 2002; Weeks, et al. 2002; Hoffmann and Weeks 2007), thus uncovering signatures of thermal adaptation. To gauge levels of mtDNA sequence variation across these populations, we used the cline end populations (Melbourne & Townsville), and estimated Fst values for each mtDNA SNP between these populations. We identified 15 SNPs in the mitochondrial genome exhibiting high (and significant) Fst values; the rest of the genome was highly conserved (Table 1, Table S1B). To probe levels of haplotype diversity, and estimate the frequencies of each haplotype within each of the source populations, we designed a custom-genotyping assay based on these 15 SNPs, and used this assay to genotype the field-collected isofemale lines (N = 312). We identified a total of 10 unique haplotypes. All haplotypes fell into one of two main haplogroups, with a total of 12 SNPs delineating the two groups (Figure 1). Both haplogroups were found to segregate across most of the 11 populations, but as a whole one haplogroup (haplogroup A) predominated in the northern sub-tropical populations while the other (haplogroup B) predominated in southern temperate populations (Figure S1). Furthermore, each of the A and B haplogroups was dominated by one major haplotype (A1 accounting for 93.3% of A haplotypes; and B1 accounting for 77.2% of B haplotypes, Figure 1). The A1 haplotype appears more closely related to other haplotypes of African origin. The ancestral origin of B1 haplotypes is, however, less clear given they are most closely related to haplotypes from other New World populations, but also a haplotype from Japan (Figure S2). Neither of these haplotypes has been previously-studied in the context of thermal selection; and indeed, no study has previously taken a clinal or intra-population approach to studying the thermal sensitivity of variation in the mtDNA genome in *D. melanogaster*. The frequency of the A1 haplotype was negatively associated with the latitude of its source population (R^2^ = 0.4847, β = –0.02881, p = 0.0273, Figure 2A), while the frequency of B1 exhibited a positive association (R^2^ = 0.5137, β = 0.02718 p = 0.0131, Figure 2B). Thus, a latitudinal cline exists for the frequencies of the A1 and B1 haplotypes along the east coast of Australia.

**Figure 1:**
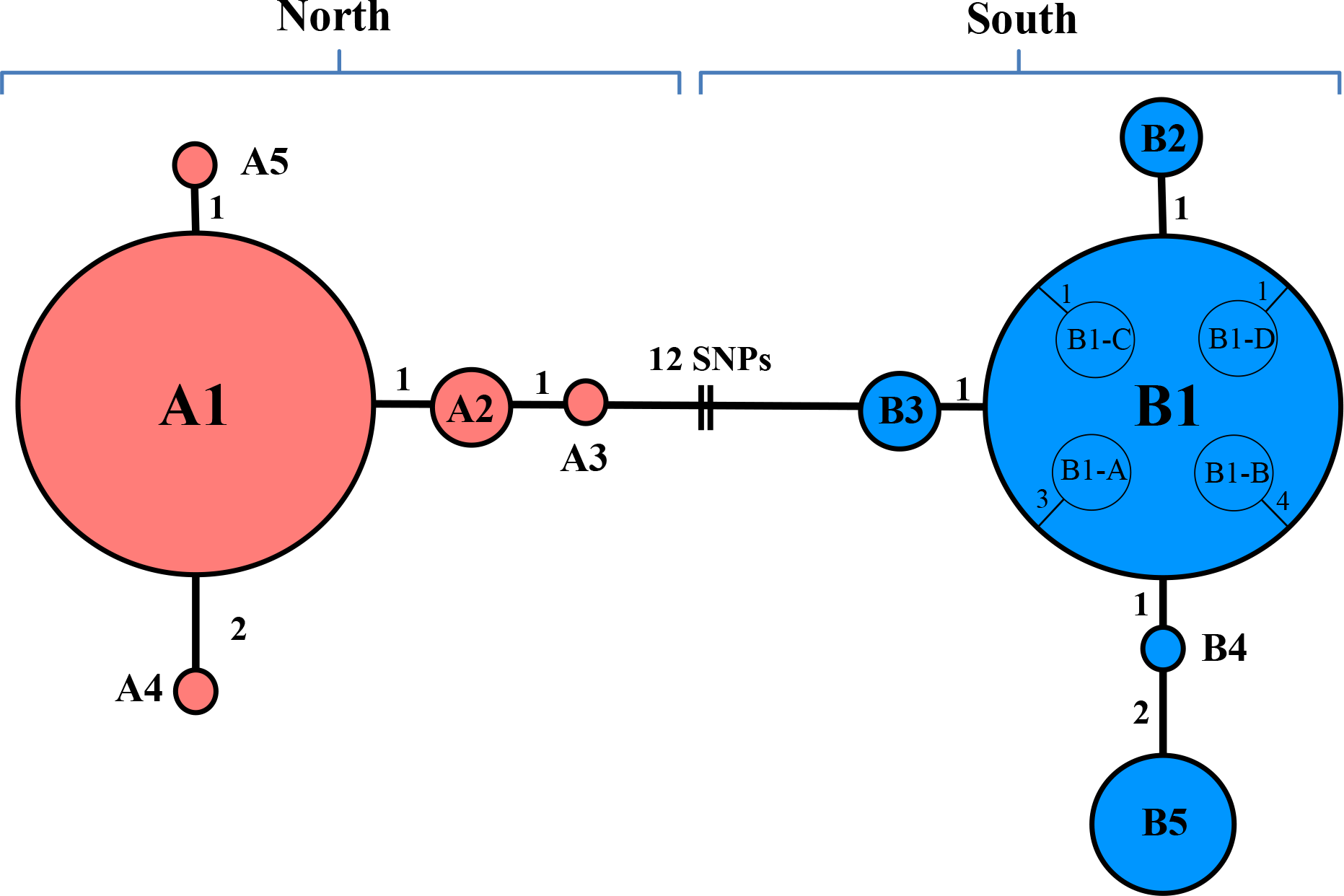
Haplotype network for mitochondrial protein-coding regions derived from genotyping of 15 SNPs. Circles indicate unique haplotypes with circle size proportional to haplotype frequency. The two different colours correspond to the two haplogroups. The A1 and B1 haplotypes were the only haplotypes present in each of the sampled populations, and are the main drivers of latitudinal association patterns (Table S1B). A1 and B1 thus contributed heavily to the frequency of each “haplogroup” per latitudinal location. In this figure, the colour red corresponds to the group of haplotypes that, when pooled together into the level of the haplogroup, is more predominant in the north of Australia (termed “North” in the figure), and blue corresponds to the pool of haplotypes that, when pooled together into the level of the haplogroup, is more predominant in the south of Australian (termed “South” in the figure; see Fig S1). Further resequencing of A1 and B1 haplotypes revealed that the B1 haplotype is comprised of at least 4 sub-haplotypes (B1-A, B1-B, B1-C, B1-D). Sub-haplotypes all share the same diagnostic 15 SNPs that delineate the B1 from the A1 haplotype, however contain 1 to 4 additional SNPs scattered throughout the mitochondrial genome (Table 2).

**Figure 2:**
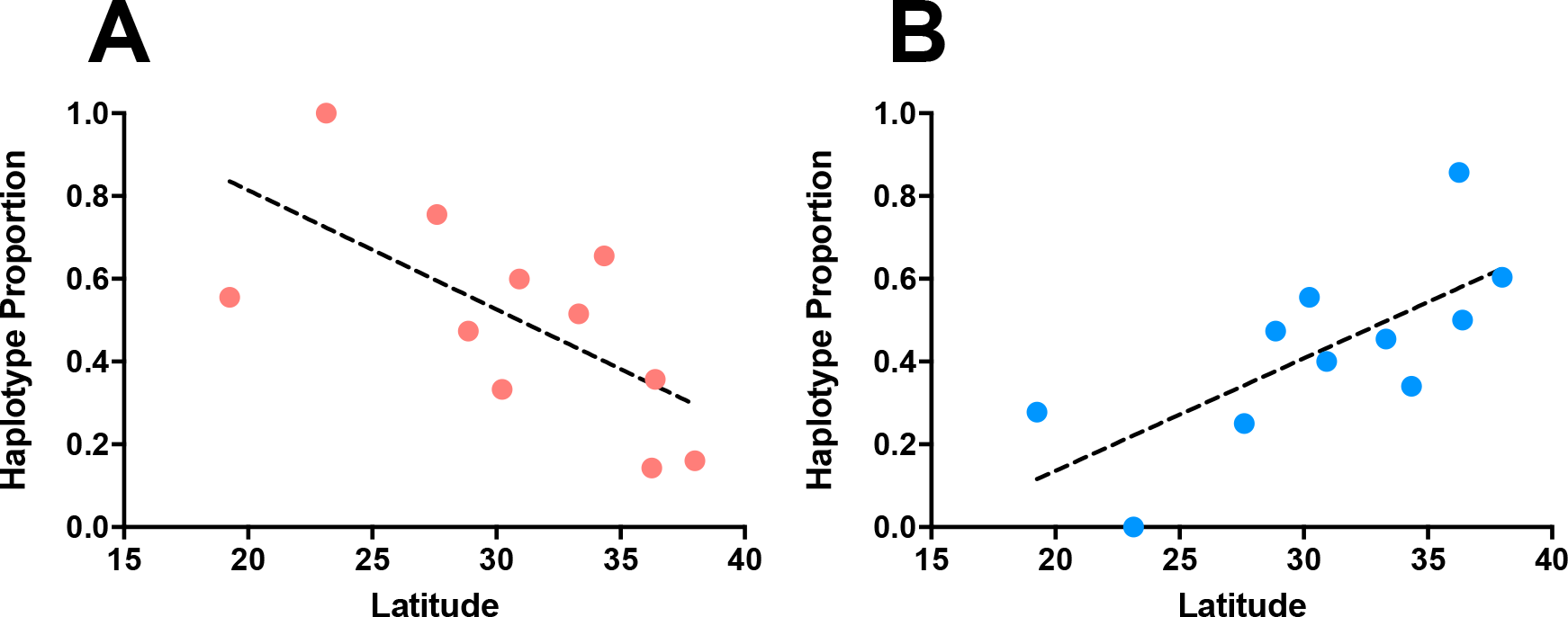
Haplotype abundance along the Australian eastern coast. **A)** Haplotype A1 (red) is predominantly found in the north of Australia, decreasing in frequency as latitude increases (R^2^ = 0.4847). **B)** Haplotype B1 (blue) is more common in the south, decreasing in frequency as latitude decreases (R^2^ = 0.5137).

**Table 1:**
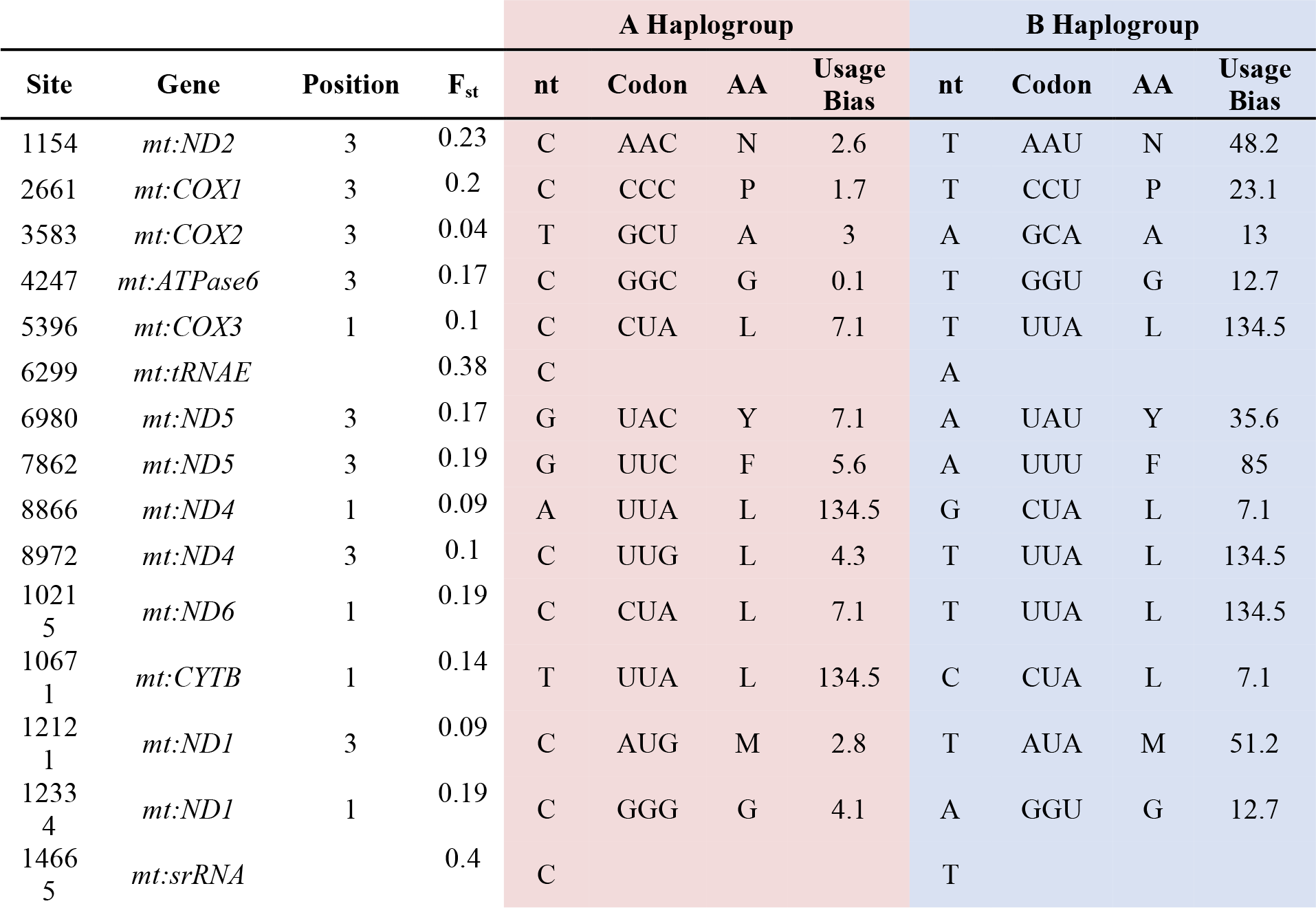
Location of all SNPs identified via next-generation sequencing of the 11 mass-bred populations. For each SNP site, we identified nucleotides that were diagnostic of the northern and southern major haplotypes. Here we list the location (Site) of the SNP, the affected gene (Gene), and the codon position (Position). Additionally, for each north and south polymorphism, we list the nucleotide (nt), the codon, amino acid (AA), and the usage bias for the specific codon. Furthermore, we provide the F_st_ values obtained from comparing the most northern and southern populations (Melbourne and Townsvile).

We next sought to experimentally assess whether the clinal associations of the A1 and B1 haplotypes are consistent with the hypothesis that these associations have been shaped by thermal selection. Mapping each of the mtDNA haplotypes to components of thermal tolerance is key to the study, given that such clinal associations could alternatively be mediated by the history of colonisation and non-adaptive demographic factors, or by other environmental variables that are likely to associate with latitudinal variation, such as humidity or dietary resources. To address this question, it was first necessary to disentangle effects attributable to mitochondrial genetic variation from those caused by segregating nuclear allelic variation, or other sources of environmental variance (Dowling, et al. 2008). We thus created eight genetic strains of flies, in which four of the strains harboured the A1 haplotype, and the other four the B1 haplotype, in an otherwise isogenic nuclear background derived from a distinct southern latitudinal population [Puerto Montt (PUE), Chile, South America]. We also ensured that all strains were free of *Wolbachia* infection, a maternally-inherited endosymbiotic bacterium, because variation in infection with different strains of *Wolbachia* would confound our capacity to map phenotypic effects to the mtDNA sequence (see Materials and Methods for more details on antibiotic treatment and *Wolbachia*-screening). Furthermore, we created these eight strains such that each haplotype was replicated across two levels (intra- and inter-latitudinal replication per haplotype), which therefore enabled us to statistically partition effects attributable to the mitochondrial haplotype from effects attributable to other sources of variation. Specifically, the haplotypes that sourced the strains were collected from each of two geographically-disjunct populations - Melbourne (37.99°S) and Brisbane (27.61°S). Because each mass-bred population was kept in independent duplicate, we ensured each duplicate contributed one A1 haplotype and one B1 haplotype to the strains (4 duplicates × 2 haplotypes = 8 strains, thus creating replication within and between latitudes, Figure S3A & B).

Once created, full protein-coding resequencing of the mitochondrial genomes of each strain revealed that those harbouring the A1 haplotype were indeed all homogeneous; characterised by a single haplotype. The strains harbouring the B1 haplotype were, however, heterogeneous (Figure 1), and could be further partitioned into four unique “sub-haplotypes” (B-1, B-2, B-3, & B-4). Each B sub-haplotype was delineated by 1 to 4 SNPs, but all shared the same pool of 12 SNPs that delineate them from the A haplogroup (Figure 1, Table 2, Figure S4). This enabled us to partition mitochondrial genetic effects over two levels – at the level of the haplotype, and the sub-haplotype. The genetic variation differentiating the A1 and B1 haplotypes was comprised of 15 synonymous SNPs in the protein-coding genes. Synonymous SNPs have traditionally been considered to be functionally silent because they do not change the amino acid sequence. However, a growing body of empirical evidence suggests that synonymous polymorphisms might routinely modify the phenotype and thus be of functional and evolutionary significance (Kimchi-Sarfaty, et al. 2007; Hurst 2011). On the other hand, the SNPs delineating the “sub-haplotypes” hubbed within the B1 haplotype, consisted of a mixture of synonymous and non-synonymous SNPs (Table 2).

**Table 2:** Location of all SNPs identified via next-generation resequencing of the mitochondrial genomes of each genetic strain, revealing that the B1 haplotype can be further partitioned into four unique “sub-haplotypes”. Below is the list comprising the origin from which each genetic strain was originally sourced (Origin), the identity of the duplicate of each population (Dup), the haplotype associated with each strain (h.type), the sub-haplotype (sub-h.type), the affected gene (Gene), whether the SNP is synonymous (S) or non-synonymous (N), the nucleotide change (nt change), the location (Site) of the SNP, and amino acid change (AA change).

| Origin | dup | h.type | sub-h.type | Gene | SNP | nt change | Site | AA change |
| --- | --- | --- | --- | --- | --- | --- | --- | --- |
| MEL | 1 | A1 | A1 |  |  |  |  |  |
| MEL | 2 | A1 | A1 |  |  |  |  |  |
| BRIS | 1 | A1 | A1 |  |  |  |  |  |
| BRIS | 2 | A1 | A1 |  |  |  |  |  |
| MEL | 1 | B1 | B1-C | *mt:COXII* | N | C → T | 3359 | P → S |
| MEL | 2 | B1 | B1-D | *mt:ND4* | S | C → T | 8033 |  |
| BRIS | 1 | B1 | B1-A | *tRNA-ASP* |  | A → C | 3892 |  |
|  |  |  |  | *mt:COXIII* | S | T → C | 4954 |  |
|  |  |  |  | *mt:ND5* | S | A → G | 7877 |  |
| BRIS | 2 | B1 | B1-B | *mt:COXI* | S | G → A | 2262 |  |
|  |  |  |  | *mt:COXII* | S | C → T | 3385 |  |
|  |  |  |  | *mt:COXIII* | N | G → A | 5162 | V → I/M |
|  |  |  |  | *mt:ND4-L* | S | A → T | 9341 |  |

Flies harbouring the A1 haplotype, which predominates in the sub-tropics, exhibited greater tolerance to an extreme heat challenge than flies harbouring the B1 haplotype (haplotype, χ^2^=6.04, p = 0.014, Table S2). However, the magnitude of these effects changed across the sexes (haplotype × sex, χ^2^=24.7, p < 0,001, Figure 3A-B, Table S2). We also uncovered sex-specific effects that mapped specifically to the level of the mtDNA sub-haplotype (sex × sub-haplotype[haplotype], χ^2^=25.04, p = <0.001, Figure 3C). This interaction was primarily attributable to the B1-D sub-haplotype, which conferred inferior heat tolerance in males, but high heat tolerance in females, relative to the other sub-haplotypes. Only one synonymous SNP, located in the *mt:ND4* gene, delineates the protein-coding region of this sub-haplotype from the other B1 sub-haplotypes (Table 2). This polymorphism is therefore a candidate SNP in conferring sex-specific outcomes in heat tolerance, although we cannot rule out the possibility that further variation within the non-coding region of the mtDNA sequence and regulatory elements (which we did not sequence) contributed to this effect. Nonetheless, the observed pattern associated with this sub-haplotype is striking in the context of a hypothesis proposed by Frank and Hurst (1996), often called *Mother’s Curse*, which proposes that maternal inheritance of the mitochondria will facilitate the accumulation of mtDNA mutations that are deleterious to males, but benign or only slightly deleterious to females (Frank and Hurst 1996; Gemmell, et al. 2004; Beekman, et al. 2014). However, while this haplotype harbours variation that causes a male-limited reduction in heat tolerance, it did not confer a detrimental effect on male capacity to tolerate cold stress (Table S3, Figure 3D). Thus, further studies are required to determine whether the male-specificity of the B1-D sub-haplotype on heat tolerance effect extends to pleiotropic effects on other life-history traits such as reproduction and longevity (Camus, et al. 2015), or whether this effect is sensitive to genotype-by-environment interactions that mediate the severity of effect in males (Mossman, et al. 2016; Wolff, et al. 2016).

**Figure 3:**
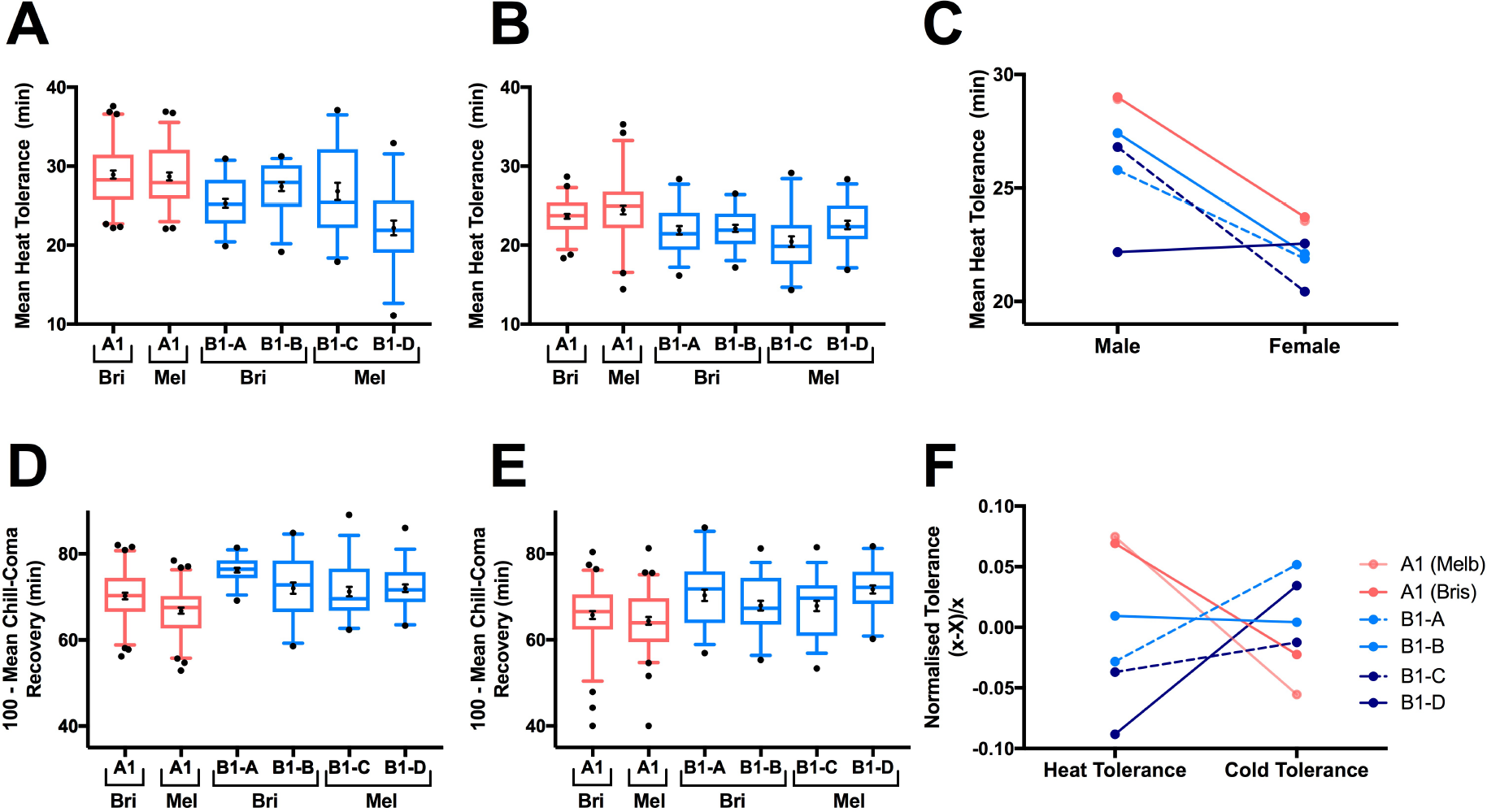
**A)** Heat tolerance (mean heat “knockdown” time ±1 S.E) of males carrying the A1 (red) and B1 (blue) haplotypes / sub-haplotypes. Means for each haplotype are shown separately according to population of origin; Bri refers to Brisbane, Mel refers to Melbourne. **B)** Heat tolerance (mean heat “knockdown” time ±1 S.E) of females carrying the A1 (red) and B1 (blue) haplotypes / sub-haplotypes. **C)** Differences in male and female heat tolerance means across mitochondrial haplotypes. **D)** Cold tolerance (100 - mean chill-coma recovery time ±1S.E) of males carrying the A1 (red) and B1 (blue) haplotypes / sub-haplotypes. **E)** Cold tolerance (100 - mean chill-coma recovery time ±1S.E) of females carrying the A1 (red) and B1 (blue) haplotypes / sub-haplotypes. **F)** Heat and cold tolerance (centred on a mean of zero and standard deviation of 1) across mitochondrial haplotypes.

Flies harbouring the B1 haplotype were superior at withstanding an extreme cold challenge, relative to their A1 counterparts (χ^2^=34.31, p < 0.001, Figure 3D-E, Table S3), but there was no significant effect traceable to the level of the sub-haplotype (Table S3). Importantly, the effects of mitochondrial haplotype on both thermal tolerance phenotypes was robust to the source of origin of the haplotypes (*i.e.*, whether they were sourced from Brisbane or Melbourne), providing clear evidence that the phenotypic effects are directly tied to the mtDNA sequence (Figure 3). Furthermore, all B1 sub-haplotypes exhibited decreased heat tolerance and increased cold tolerance when compared to the A1 haplotype, providing support for the suggestion that the differences in thermal tolerance observed between northern and southern haplogroups are mapped to the 15 SNPs that delineate the A1 and B1 haplotypes (or to cryptic variation in the non-coding region that we did not genotype), rather than the SNPs that delineate the different sub-haplotypes hubbed within B1 (Table S2, Table S3, and Figure 3F). Alternatively, it is possible that the unique SNPs that delineate the B1 sub-haplotypes drive the bulk of the differences in thermal response between the A1 and B1 haplotypes, and represent cases of parallel evolution for thermal tolerance brought about by different underlying SNPs (Arendt and Reznick 2008). While we cannot definitively disentangle these possibilities, we note that the polymorphisms that delineate the A1 from the B1 sub-haplotypes only include non-synonymous SNPs in two of four cases. Thus, while we are unable to definitively ascertain whether differences in the A1 and B1 thermal responses are underpinned primarily by the 15 shared SNPs that separate all B1 from A1 haplotypes, by other cryptic regulatory variation in the non-coding region, or by the unique SNPs that delineate the B1 sub-haplotypes, our results suggest that SNPs that do not change the amino acid sequence are likely to be responsible for this thermal divergence in at least two cases.

We then examined whether the thermal tolerance phenotypes might be mediated by patterns of differential gene expression of protein-coding mtDNA genes, copy number variation in mtDNA, or codon usage bias across the A1 and B1 haplotypes. While we did not detect differences in mtDNA copy number between A1 and B1 haplotypes (Table S5), we did detect differences in mitochondrial gene expression. Specifically, we extracted RNA of females of the A1 and B1 haplotypes, and examined expression patterns in five genes involved in complex I and complex IV of the electron transport chain (complex I: *mt:ND4*, *mt:ND5,* complex IV: *mt:COXI*, *mt:COXII*, *mt:COXIII*). Emerging evidence suggests that genetic variation within complex I genes (both mitochondrial and nuclear) might contribute disproportionately to trajectories of mitonuclear, and ultimately, life history evolution (Camus, et al. 2015; Garvin, et al. 2015; Morales, et al. 2015). Complex IV, on the other hand, harbours genes with the lowest levels of *dN/dS*, indicative of greater selective constraints on these mitochondrial genes across taxonomically-diverse organisms (Nabholz, et al. 2013). Accordingly, we found that strains harbouring the B1 haplotype exhibited higher gene expression for the complex I genes *mt:ND4* and *mt:ND5*, which belong to the same transcriptional unit (Torres, et al. 2009), than strains with the A1 haplotype (haplotype × gene < 0.001, Figure 4, Table S4). We note we conducted these analyses in females only, since a study of mitochondrial gene expression across a global sample of mtDNA haplotypes had previously indicated that mtDNA haplotypes affect the expression of protein-coding mtDNA genes, but that these haplotype-specific effects are consistent across the sexes (Camus et al 2015). Future work could, however, examine whether these patterns of gene expression across the A1 and B1 haplotypes are upheld in males, and whether the differences in *mt:ND4* and *mt:ND5* expression observed across haplotypes, extend further to differences at the level of the individual B1 sub-haplotypes. We note that all of the SNPs located in the *mt:ND4* and *mt:ND5* genes, which delineate A1 from B1 haplotypes, are synonymous (Table 1). This observation is interesting in light of a recent report that found that patterns of expression of *mt:ND5* and *mt:CYTB* genes in *D. melanogaster* mapped to candidate SNPs that lay directly within the affected genes, and which presumably exerted their effects via post-transcriptional modification of RNA, potentially altering the stability of the transcripts (Camus, et al. 2015). This is also consistent with reports showing that coding variants in both the nuclear and mitochondrial genome affect gene expression patterns in humans (Birnbaum, et al. 2012; Cohen, et al. 2016), and recent evidence demonstrating that transcription regulators specifically bind to the human mtDNA coding region to regulate transcription (She, et al. 2011; Blumberg, et al. 2014). In combination, these studies suggest that the synonymous SNPs delineating the A1 and B1 haplotypes could be involved in regulating transcription of these genes, via mito-nuclear interactions involving nuclear-encoded transcription regulators.

**Figure 4:**
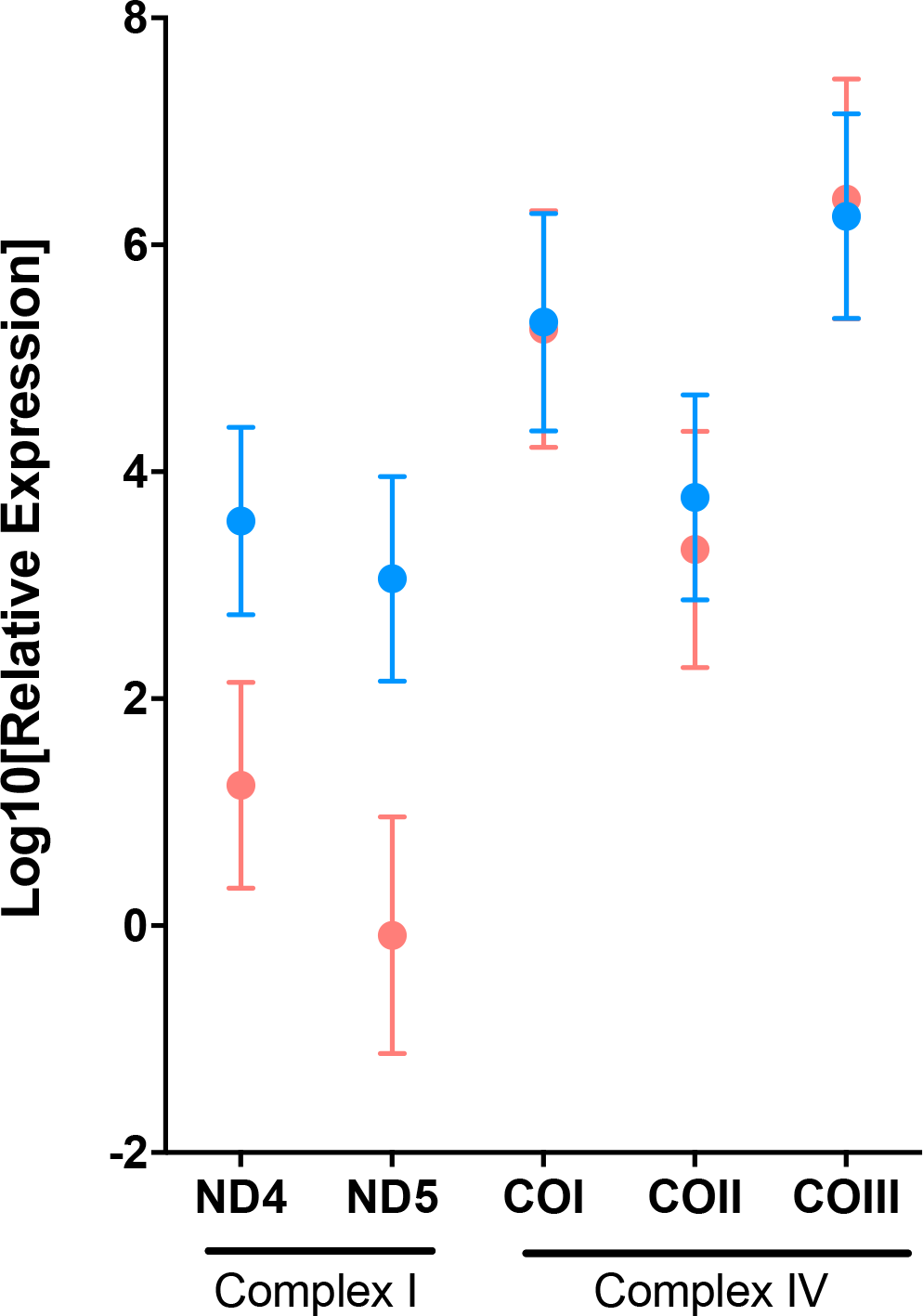
Least-squares means (± 1 S.E.) of female gene expression across A1 (red) and B1 A-D combined (blue) haplotypes for the *mt:ND4* and *mt:ND5* genes (OXPHOS complex 1) and *mt:COI*, *mt:COII*, *mt:COII* genes (OXPHOS complex IV). *mt:COI*, *mt:COII*, *mt:COII* all belong to one transcriptional unit and encode subunits of complex IV, whilst *mt:ND4* and *mt:ND5* are members of a second transcriptional unit and encode subunits of complex 1 of the mitochondrial electron transport chain. Least-square means for all plots were derived from the multilevel models, which take into account mtDNA copy number as a covariate (Table S4)

Evidence is also mounting that variation in patterns of genomic DNA base composition (GC content (Šmarda, et al. 2014), as well as variation in codon usage bias across DNA sequences (Sharp, et al. 1995) can be shaped under natural selection. For example, in bacteria and metazoans, higher levels of GC base pairs have been associated with the thermal environment, with the GC base pair associated with higher thermal stability (Bernardi 2007). In bacteria, this correlates with greater tolerance of higher temperatures (Musto, et al. 2004). In the green alga *Chlamydomonas*, experimental alteration of mitochondrial codons drastically changes translational efficiency, suggesting that mitochondrial codon usage has been optimised for translation of mitochondrial products (Salinas, et al. 2012). In our study, the A1 and B1 haplotypes differ by 15 synonymous SNPs that are evenly distributed across the mitochondrial protein-coding region, with most protein-coding genes harbouring at least one SNP site. SNPs of the A1 haplotype show a high GC bias, with 80% of the SNPs represented by a guanine or cytosine, and conversely those of B1 reveal a GC content of only 20% (Table S6A, Fishers exact test, p = 0.001). Thus, the A1 haplotype, which confers higher tolerance to an extreme heat challenge, has a higher GC content; concordant with previous observations in bacteria and metazoans suggesting higher thermal stability of the GC base pair relative to AT (Bernardi 2007). Additionally, the SNPs delineating the A1 haplotype change the codon bias and produce rarer codons (Table S6B, Fishers exact test p, = 0.002). These findings suggest that GC content and codon bias may play a role in the observed haplotype effects on gene expression of *mt:ND4* and *mt:ND5*, with ultimate upstream effects on thermal tolerance.

By harnessing an experimental genomic approach applied to the mitochondrial genome, within a clinal framework, we have documented latitudinal patterns in standing mtDNA haplotypes, and provided experimental evidence that these patterns are linked to the capacity of these haplotypes to tolerate thermal stress. While these results are consistent with the suggestion that the clinal patterns of mtDNA variation are likely to have been shaped at least in part by thermal selection, it is difficult to fully resolve the relative influence of thermal selection from history of colonisation and other demographic factors, given that the Australian east coast is thought to have been subjected to recurrent colonisation events from flies of disparate origins over the past 150 years. This, of course, is a caveat that is not unique to our study on mtDNA variation, but extends to all studies of clinal variation of New World populations of *D. melanogaster*. We thus point out that these haplotypes might have been pre-adapted to tropical and temperate conditions of Africa and Eurasia prior to their introductions into Australia, and that the relevant mitochondrial variation under selection along the Australian cline is likely to have already existed upon the arrival of these haplotypes into Australia. Accordingly, the Australian distribution of mtDNA haplotypes is likely to have been shaped both by the history of colonization, followed by the subsequent and ongoing action of thermal selection in spatially-sorting the haplotypes along the latitudinal cline. Our study suggests that further research into the mitochondrial climatic adaptation hypothesis is warranted. In particular, we suggest our conclusions can be tested by future studies that utilise other established latitudinal clines, in *D. melanogaster* and in other species, to determine whether mtDNA haplotypes show similar associations to latitude as revealed in the Australian cline, and to determine whether the mtDNA haplotypes involved exhibit thermal sensitivities that concord with the clinal patterns.

In conclusion, our results provide support for the hypothesis that standing genetic variation within the mitochondrial genome has been shaped, in part, by natural selection imposed by thermal stress. We also presented evidence that SNPs found within the mtDNA, and which do not change the amino acid sequence, contribute to the regulation of phenotypic responses to thermal stress in *D. melanogaster*. This thus suggests a role for a set of SNPs that were traditionally thought to evolve under neutrality (Kimchi-Sarfaty, et al. 2007; Hurst 2011), within a genome that was likewise traditionally thought to be devoid of phenotype-changing genetic variation (Ballard and Rand 2005; Dowling, et al. 2008), in the dynamics of thermal adaptation. Secondly, and more broadly, our results add to an emerging body of research in *Drosophila* (Sorensen, et al. 2007; Chen, et al. 2012; Lavington, et al. 2014; Cogni, et al. 2015) and other metazoans (Porcelli, et al. 2015), which highlights metabolic genes (including those targeted to the mitochondria) as important substrates on which thermal selection is likely to act to shape adaptive evolutionary responses. Several studies have now reported variation in allele frequencies, or expression patterns, of nuclear-encoded metabolic genes along latitudinal clines (Chen, et al. 2012; Lavington, et al. 2014; Cogni, et al. 2015), or across replicated laboratory populations that have evolved under differing thermal regimes (Sorensen, et al. 2007), in *Drosophila melanogaster*. These studies, however, did not screen for involvement of the evolutionary-conserved mitochondrial genes. The function of key metabolic enzymes, however, relies on close coordination between mitochondrial and nuclear genomes (Rand, et al. 2004; Levin, et al. 2014; Wolff, et al. 2014; Quiros, et al. 2016). This point, when reconciled with the emerging studies, would suggest that genetic interactions between the mitochondrial and nuclear genomes could represent key mediators of evolutionary adaptation of the metabolic machinery under thermal stress.

## Materials and Methods

### Field Collection, Isofemale Line Establishment and Maintenance

Populations of *Drosophila melanogaste*r were sampled from the east coast of Australia during March-April 2012 from 11 locations. The population names (latitude and longitude) are: Townsville (19.26,146.79), Rockhampton (23.15, 150.72), Brisbane (27.61, 153.30), Ballina (28.87,153.44), Coffs Harbour (30.23, 153.15), Port Macquarie (30.93, 152.90), Wollongong (34.34, 150.91), Narooma (36.25, 150.14), Gosford (33.31, 151.20), Bermagui (36.40, 150.06), Melbourne (37.99, 145.27). Samples were collected as close to sea level as possible to avoid altitudinal differences between the populations (Collinge, et al. 2006). Individual field-inseminated females were isolated into individual vials in the laboratory to initiate independent isofemale lines. At least twenty isofemale lines were generated for each population. Each line was treated with tetracycline to eliminate cytoplasmic endosymbionts, such as *Wolbachia* (Clancy and Hoffmann 1998), and tested using *Wolbachia*-specific primers (O’Neill, et al. 1992). We further verified infection status when analysing next-generation sequencing data by confirming that none of our obtained reads mapped to the *Wolbachia* genome (NC_002978).

Three generations after the isofemale lines were established in the lab, one mass-bred population was created from the isofemale lines of each latitudinal location (11 locations). Specifically, the populations were established by combining 25 virgin males and 25 virgin females from each of 20 randomly-selected isofemale lines per latitudinal location. The following generation, each population was divided into two duplicates (11 populations × 2 duplicates), which were kept separately from this point onward. A small sample of flies (~20–50 individuals) from each isofemale line was also collected at this time, and placed at –20°C for sequencing and genotyping. Mass-bred populations were kept at 25°C under a 12:12h light:dark cycle. Genetic variation was maintained within each duplicate population by rearing flies across two bottles on potato-dextrose-agar food medium, with densities of approximately 300 flies per bottle. Every generation, newly-emerged flies from each duplicate were collected from both bottles and then randomly redistributed into two new fresh bottles.

### Next Generation Sequencing and SNP Genotyping

To identify regions of variation between the 11 populations, we first used pooled samples of 100 individuals (both males and females) from each population and used next generation sequencing to obtain full mitochondrial genomes. DNA samples were enriched for mitochondrial DNA to obtain the best coverage possible. This process was achieved by using *Wizard SV* Miniprep Purification Kit (Promega, Madison, WI) for DNA extraction, which captures circular DNA. Enriched DNA samples were made into 200bp paired-end libraries and sequenced using the Illumina GAIIx platform (Micromon, Monash University, Australia). Reads were aligned to the *Drosophila melanogaster* mitochondrial reference genome (NCBI reference sequence: NC_001709.1) using *Geneious* (Kearse, et al. 2012), generating mitochondrial protein coding regions for each of the 11 latitudinal locations. Given the high A-T richness of the mitochondrial genome of *Drosophila melanogaster* it is extremely difficult to map reads to the control region (D-loop). The D-loop is a 5kb repetitive region with A-T richness of over 90%, making this region extremely difficult to accurately map reads (Tsujino, et al. 2002). Although our level of coverage was more than sufficient to examine the protein-coding region, we were not able to accurately map sufficient reads to the D-loop.

We first aimed to identify SNPs in the mitochondrial genome that had high levels of genetic differentiation between locally extreme populations. For this we used poolSeq data from the cline extremes (Melbourne and Townsville) and calculated FST values for individual SNPs of the mitochondrial genome using *Popoolation2* (Kofler, et al. 2011). To obtain allele frequencies from each population, SNP sites with high Fst were used as markers. DNA from each isofemale line was extracted using the *Gentera Puregene Cell and Tissue Kit* (Qiagen, Hilden, Germany). Even though each mass-bred population was created using 20 randomly-chosen isofemale lines, we genotyped all isofemale lines collected from each latitudinal location. A custom SNP genotyping assay was developed (Geneworks, Thebarton, Australia) for the 15 SNPs identified via mass sequencing, and genotyping was performed by Geneworks (Thebarton, Australia) on a SEQUENOM MassARRAY platform (Agena Bioscience, San Diego, CA). This genotyping revealed the presence of northern-predominant (*i.e.,* predominating in northern latitude populations) and southern-predominant (*i.e.*, predominating in southern latitude populations) haplogroups, with each haplogroup characterised by one major haplotype (A1 and B1).

We also relied on published genomic datasets to screen for signatures of latitudinal variation in mtDNA SNPs within established latitudinal clines of *D. melanogaster* along the east coast of North America, and Africa. These analyses, and interpretations, are presented in the Supplementary Information.

### Creation of Mitochondrial Strains from Mass-bred Populations

We created “introgression strains” from each of the population duplicates (11 latitudes × 2 population duplicates = 11 introgression strains × 2 duplicates), by introgressing the pool of mtDNA variants of each population duplicate into a standard and isogenic nuclear background originally sourced from Puerto Montt (PUE), Chile (41.46°S, 72.93°W) (Calboli, et al. 2003), which had been created via 20 generations of full-sibling matings. We chose this background primarily because it was from a distinct southern-hemisphere, and is very unlikely to have shared a recent coevolutionary history with either of the A1 and B1 haplotypes (see Fig S4), which might have inadvertently favoured one or other of the mtDNA haplotypes in our phenotypic assays of thermal tolerance. To initiate each strain, 100 virgin female flies were sampled from each population duplicate and crossed to 120 males from the PUE strain. Then, for 20 sequential generations, 100 daughters were collected per strain and backcrossed to 120 PUE males. This crossing scheme aimed to maintain the pool of segregating mitochondrial haplotypes within each population, while translocating them alongside that of an isogenic nuclear background, to enable partitioning of mitochondrial genetic effects from cryptic variance tied to the nuclear genome (Figure S3A). In order to prevent mitochondrial contamination from the Puerto Montt (PUE) line, all lines were tested every 5 generations during the introgression regime, to ensure there were no instances of contamination of the lines (by rogue females of the PUE strain) by using qRT-PCR melt curve analysis that would detect PUE-specific mtDNA SNPs.

We then created a new set of isofemale lines from each of the introgression strain duplicates, and re-genotyped females of each line using the custom SNP genotyping assay described above (Geneworks, Thebarton, Australia). From the genotyping results, we were able to identify female lineages that carried individual haplotypes (A1 [northern] or B1 [southern]), and using this information we then selected one isofemale line carrying the A1 haplotype and one isofemale line carrying the B1 haplotype, from each of the two independent population duplicates from two (Brisbane, Melbourne) of the 11 latitudinal locations (Figure S3B). We continued to backcross virgin females of each isofemale line to males of the isogenic PUE line for a further seven generations. We acknowledge that in the presence of strong mito-nuclear coevolution, such a backcrossing approach could in theory fail to disrupt essential allelic pairings spanning mitochondrial and nuclear genotypes, meaning that a few nuclear alleles that are essential to maintaining mito-nuclear compatibility might remain, even following 27 generations of backcrossing. While a theoretical possibility, this seems unlikely from a population genetic perspective (Eyre-Walker 2017); particularly in our study, given that the A1 and B1 haplotypes under introgression here co-occur within the same panmictic populations, and differ only by a small number of SNPs don’t change the amino acid sequence (Table 1), and given that we have never previously come across combinations of mito-nuclear genotypes in *D. melanogaster* (even at the inter-population scale) that incur complete inviability in females (the sex that transmits the mtDNA) or juveniles. Prior to this step, the PUE line had been propagated via a protocol of mating between one full-sibling pair for five generations, to remove any genetic variation that had accumulated within this nuclear background during the course of the introgressions described above. We chose to use isofemale lines from Brisbane (latitude: 27.61°S) and Melbourne (latitude: 37.99°S) because they are geographically-disjunct, and because re-genotyping confirmed that both A1 and B1 haplotypes were segregating in each of the introgression strain duplicates following the 20 generations of introgression. Following this process, each of the A1 and B1 haplotypes was represented across four independent genetic strains each, at two levels of replication; an intra-latitudinal (between the two population duplicates of a given latitude) and an inter-latitudinal (between two latitudes, Brisbane and Melbourne) replicate (Figure S3B).

We then re-sequenced these strains, and obtained full complete mitochondrial genomes for all eight mitochondrial strains, again using the next generation sequencing approach described above. Resequencing results revealed that haplotype A1 was isogenic across all four A1 strains, while we found that the B1 strains could be delineated into four unique sub-haplotypes that were nested within the B1 haplotype. These four southern sub-haplotypes all shared the known SNPs that delineate the north and south haplogroups (and the A1 and B1 haplotypes), however they each carried between one and three additional SNPs (Table 2).

### Extreme Heat Challenge

Tolerance to an extreme heat challenge was measured for 120 flies of each sex from each mitochondrial strain (Hoffmann, et al. 2002). Flies were placed in individual 5mL water-tight glass vials and subsequently exposed to a 39°C heat challenge, by immersion of the glass vials in a preheated circulating water bath. Heat “knock-down” time was recorded as the time taken for each individual fly to become immobilized (in a coma-like state) at 39°C (Williams, et al. 2012). This experiment was conducted over two trials within the same generation. Each trial of the experiment consisted of a fully-balanced replicate of the experimental units (*i.e.*, equal numbers of flies of each sex × mitochondrial strain), separated in time by 2 hours within the same day. The position of flies of each experimental unit was randomized within each trial of the experiment. The assay was conducted blind to the genotype or sex of the fly.

### Extreme Cold Challenge

This assay measures the amount of time it takes a fly to regain consciousness and stand on all legs after succumbing to a cold-induced coma (Hoffmann, et al. 2002). In each trial of the assay, 40 flies from each experimental unit (N = 640) were placed individually in 1.7mL microtubes. These tubes were then submerged in a water bath set to 0°C (comprised of water and engine coolant) for 4 h, to place flies into coma. At 4 h, all microtubes were removed from the bath, and laid out on a bench at 25°C, and the time taken (seconds) for each fly to regain consciousness and stand upright was recorded. The assay was conducted blind to the genotype or sex of the fly.

### Statistical Analyses of Thermal Tolerance Data

We used separate multilevel linear mixed models to test the effects of mtDNA haplotype and sub-haplotype on responses to each of the heat and cold challenges. The response variable for the heat challenge assay was the time taken to fall into coma, while the response variable for the cold challenge assay was the time taken to wake from coma. Fixed effects were the identity of the mtDNA haplotype (A1, B1), the sub-haplotype nested within haplotype (A1, B1-A, B1-B, B1-C, B1-D), sex and their interactions. This analysis assumes the effect of the SNPs separating the A1 and the B1 haplotypes, and those that separate the B1 sub-haplotypes, are hierarchical and can be statistically partitioned (*i.e.*, any significant ‘haplotype’ effects in the model can be mapped to the 15 SNPs that separate the A1 and B1 haplotypes, while significant ‘sub-haplotype’ effects are mapped to the unique SNPs that separate the four B1 sub-haplotypes). We, however, acknowledge the alternative possibility that the unique SNPs separating B1 sub-haplotypes could in theory underpin the differences between the A1 and B1 thermal responses, if such SNPs have accumulated under parallel evolution (Arendt and Reznick 2008). Random effects described the biological structure of the mitochondrial strains; there were two tiers of replication – with each haplotype replicated across two “duplicates” within each of two latitudinal “populations”. Thus, duplicate nested within population was included as a random effect, as well as other known and random environmental sources of variance (the trial identity, and the identity of the person scoring the response variable [2 people]).

Parameter estimates were calculated using restricted maximum likelihood algorithm in the *lme4* package of R (Bates 2012). The fitted model was evaluated by simplifying a full model, by sequentially removing terms that did not change (at α = 0.05) the deviance of the model, starting with the highest order interactions, using log-likelihood ratio tests to assess the change in deviance in the reduced model relative to the previous model (Fox 2002).

### Haplotype Network, Divergence, Codon Bias and RNA analysis

Relationships among haplotypes were visualized on a median-joining network (Bandelt, et al. 1999) and constructed in the software NETWORK version 4.6.1.2 (www.fluxus-engineering.com).

We obtained divergence estimates between A1 and B1 haplotypes using *Geneious* (Kearse, et al. 2012) and *MEGA6* (Tamura, et al. 2013). Using *Geneious*, divergence was calculated by examining the *%identity function* and subtracted that value from 100 to derive the percentage divergence. In *MEGA6*, we performed a pairwise distance comparison using a maximum composite likelihood model. Both methods gave concordant estimates of divergence (divergence = 0.001%)

We obtained *Drosophila melanogaster* mitochondrial codon usage bias values from the Codon Usage Database(Nakamura, et al. 2000). For both haplotypes, each SNP site was given the title “preferred” or “unpreferred” based on the codon usage bias score. Results were then analysed as a 2 × 2 contingency table using Fishers exact tests (Table S6A & B).

Sequence polymorphisms in structural RNAs between the two haplogroups were analyzed using tRNAScan (Schattner, et al. 2005) for tRNAE and ExpaRNA (Smith, et al. 2010) for the polymorphism present in the small ribosomal subunit. Secondary structures are presented in the Figure S5 and Figure S6.

### Total RNA/DNA Extraction and cDNA Synthesis

For RNA extractions, we placed single female flies from each A1 and B1 strain into a microtube. We thus combined source population and duplicate into one sample. Each extraction was performed in triplicate, thus resulting in three microtubes with flies possessing the A1 haplotype and three microtubes with flies harbouring the B1 haplotype. In the case of the A1 haplotype all flies harboured the same haplotype (although originating from different rearing vials), whereas for the B1 haplotype each biological replicate was formed by combining a single fly from each sub-haplogroup into a microtube.

We then performed a coupled RNA and DNA extraction as per the supplier’s protocols using *TRIzol® Reagent* (Thermo Fisher Scientific, Waltham, MA) to first create a phase separation of RNA and DNA from which the total RNA was then purified using a *HighPure RNA extraction kit* (Roche Applied Science, Penzberg, Germany). In this manner, both the DNA and RNA was independently separated and stored from the one sample. The separated nucleic acids (~100µL of each sample extracted) were quantified by Nanodrop UV/Vis spectrophotometry (Thermo Fisher Scientific, Waltham, MA) and the purity of total RNA was confirmed using the A_260_/A_280_ ratio with expected values between 1.8 and 2.0. The integrity of both the RNA and DNA was assessed by electrophoresis (1% TBE agarose gel).

The cDNA was synthesized from 1*µ*g of RNA using the *Transcriptor First Strand cDNA Synthesis* Kit (Roche Applied Science, Penzberg, Germany) and a mixture of random hexamers and oligodT primers to capture mitochondrial transcripts both in the transitory polycistronic stage and as individual polyadenylated single transcripts (Clayton 2000).

### mtDNA Copy Number Quantification

mtDNA copy number was measured for each DNA extraction performed (see *Total RNA/DNA Extraction and cDNA Synthesis*). MtDNA copy number was calculated relative to a single copy gene in the nuclear genome (Correa, et al. 2012). Copy number was determined using quantitative real-time PCR of a 113 bp region of the large ribosomal subunit (CR34094, FBgn0013686). No nuclear copies of this gene are found in the *Drosophila melanogaster* genome. Similarly, nuclear DNA was quantified by amplifying a 135 bp region of the single-copy (Aoyagi and Wassarman 2000) subunit of the RNA polymerase II gene (CG1554, FBgn0003277). The copy number was then determined as the relative abundance of the mtDNA to nuclear DNA ratio and thus reflects the average number of mtDNA copies per cell.

### Gene expression quantification

Five of the thirteen total mitochondrial protein-coding genes were amplified to quantify gene expression levels. Quantified genes were: *mt:COI, mt:COII, mt:COIII, mt:ND4, and mt:ND5*. Gene expression of each biological replicate (three biological replicates per haplotype) was measured using quantitative real time (qRT)-PCR (Lightcycler 480 – Roche Applied Science, Penzberg, Germany). Reactions were performed in duplicate (technical duplicates) using a *SYBRGreen I Mastermix* (Roche Applied Science, Penzberg, Germany), whereby each well contained 5µl of SYBR buffer, 4µl of 2.5µM primer mix and 1µl of diluted cDNA. The following amplification regime used was: 90°C (10s), 60°C (10s), 72°C (10s) for 45 cycles, followed by a melt curve analysis to verify the specificity of the primer pair.

The *Bestkeeper©* software (Pfaffl, et al. 2004) was used to select nuclear housekeeping genes (HKGs) for quality assessment. Three suitable HKGs were chosen: succinate dehydrogenase A (CG17246), 14–3–3ε (CG31196), and an unknown protein-coding gene (CG7277). All three genes had similar expression levels with high correlation coefficients (>0.8) against each other. For each experimental sample, the expression values of the mitochondrial target genes were standardized as follows:

The cycle threshold was calculated using the gene of interest (GOI) and the geometric mean of the three housekeeping genes (GEOM):

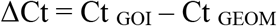

The cycle thresholds were then used to calculate the relative gene expression for each experimental sample in relation to the housekeeping genes.

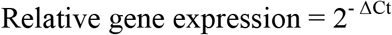

Gene expression levels of all five mitochondrial genes were obtained by determining the ΔCt per sample, measured at the maximum acceleration of fluorescence, using the Second Derivative Maximum Method (Rasmussen 2001) in the *Lightcycler Software V1.5.0* (Roche Applied Science, Penzberg, Germany). When the ΔCt values between two technical duplicates for each sample fell within 0.5 units of each other, then the mean gene expression estimates were pooled to form a single data point (Bustin, et al. 2009).

### Statistical Analysis of Gene Expression Data and Copy Number Variation

We fitted linear models, in which mitochondrial copy number and gene expression data were modelled separately as response variables. Mitochondrial haplotype (A1, B1), and gene identity were modelled as fixed effects. Mitochondrial copy number values were added as a fixed covariate to the analysis of gene expression, and F statistics and associated probabilities estimated using a Type III sums-of-squares tests in the *car* package (Fox 2002) in *R* (R Development Core Team 2009). Mitochondrial copy number variation was modelled with haplotype (A1 and B1) as a factor.

## Acknowledgements

We would like to thank Fiona Cockerell, Allannah Clemson, Winston Yee and Belinda Williams for assistance with the heat and cold tolerance assays. We thank Vanessa Kellerman and Winston Yee for fly collection. This research was supported by funding from the Hermon Slade Foundation (HSF 15/2), Australian Research Council (DP1092897, FT160100022), and Monash University Research Fellowship to DKD. CMS was funded by an ARC fellowship and the Science and Industry Endowment Fund. During part of the writing/analyses MFC was supported by a Monash Postgraduate Publications Award and the European Research Council under the Marie Skłodowska-Curie Actions (#708362).

## References

Adrion JR, Hahn MW, Cooper BS. 2015. Revisiting classic clines in Drosophila melanogaster in the age of genomics. Trends in Genetics 31:434–444.

Aoyagi N, Wassarman DA. 2000. Genes encoding Drosophila melanogaster RNA polymerase II general transcription factors: diversity in TFIIA and TFIID components contributes to gene-specific transcriptional regulation. Journal of Cell Biology 150:F45–50.

Arendt J, Reznick D. 2008. Convergence and parallelism reconsidered: what have we learned about the genetics of adaptation? Trends in Ecology & Evolution 23:26–32.

Arnqvist G, Dowling DK, Eady P, Gay L, Tregenza T, Tuda M, Hosken DJ. 2010. Genetic Architecture of Metabolic Rate: Environment Specific Epistasis between Mitochondrial and Nuclear Genes in an Insect. Evolution 64:3354–3363.

Ballard JW, Kreitman M. 1994. Unraveling selection in the mitochondrial genome of Drosophila. Genetics 138:757–772.

Ballard JWO, Rand DM. 2005. The population biology of mitochondrial DNA and its phylogenetic implications. Annual Review of Ecology Evolution and Systematics 36:621–642.

Ballard JWO, Whitlock MC. 2004. The incomplete natural history of mitochondria. Molecular Ecology 13:729–744.

Balloux F, Handley LJL, Jombart T, Liu H, Manica A. 2009. Climate shaped the worldwide distribution of human mitochondrial DNA sequence variation. Proceedings of the Royal Society Biological Sciences Series B 276:3447–3455.

Bandelt HJ, Forster P, Rohl A. 1999. Median-joining networks for inferring intraspecific phylogenies. Molecular Biology and Evolution 16:37–48.

Bates D, Maechler, M., Bolkler, B. 2012. lme4: Linear mixed-effects models using S4 classes. R package version 0.999999-0. http://cran.r-project.org/package=lme4.

Bazin E, Glémin S, Galtier N. 2006. Population Size Does Not Influence Mitochondrial Genetic Diversity in Animals. Science 312:570–572.

Beekman M, Dowling DK, Aanen DK. 2014. The costs of being male: are there sex–specific effects of uniparental mitochondrial inheritance? Philosophical Transactions of the Royal Society of London B Biological Sciences 369:20130440.

Bergland AO, Tobler R, Gonzalez J, Schmidt P, Petrov D. 2016. Secondary contact and local adaptation contribute to genome-wide patterns of clinal variation in Drosophila melanogaster. Molecular Ecology 25:1157–1174.

Bernardi G. 2007. The neoselectionist theory of genome evolution. Proceedings of the National Academy of Sciences of the United States of America 104:8385–8390.

Birnbaum RY, Clowney EJ, Agamy O, Kim MJ, Zhao JJ, Yamanaka T, Pappalardo Z, Clarke SL, Wenger AM, Nguyen L, et al. 2012. Coding exons function as tissue-specific enhancers of nearby genes. Genome Research 22:1059–1068.

Blumberg A, Sailaja BS, Kundaje A, Levin L, Dadon S, Shmorak S, Shaulian E, Meshorer E, Mishmar D. 2014. Transcription Factors Bind Negatively Selected Sites within Human mtDNA Genes. Genome Biology and Evolution 6:2634–2646.

Burton RS, Pereira RJ, Barreto FS. 2013. Cytonuclear Genomic Interactions and Hybrid Breakdown. Annual Review of Ecology, Evolution, and Systematics, Vol 44 44:281–302.

Bustin SA, Benes V, Garson JA, Hellemans J, Huggett J, Kubista M, Mueller R, Nolan T, Pfaffl MW, Shipley GL, et al. 2009. The MIQE Guidelines: Minimum Information for Publication of Quantitative Real-Time PCR Experiments. Clinical Chemistry 55:611–622.

Calboli FC, Kennington WJ, Partridge L. 2003. QTL mapping reveals a striking coincidence in the positions of genomic regions associated with adaptive variation in body size in parallel clines of *Drosophila melanogaster* on different continents. Evolution 57:2653–2658.

Camus MF, Wolf JB, Morrow EH, Dowling DK. 2015. Single Nucleotides in the mtDNA Sequence Modify Mitochondrial Molecular Function and Are Associated with Sex–Specific Effects on Fertility and Aging. Current Biology 25:2717–2722.

Chen Y, Lee SF, Blanc E, Reuter C, Wertheim B, Martinez–Diaz P, Hoffmann AA, Partridge L. 2012. Genome-wide transcription analysis of clinal genetic variation in Drosophila. PLoS One 7:13.

Cheviron ZA, Brumfield RT. 2009. Migration–Selection Balance and Local Adaptation of Mitochondrial Haplotypes in Rufous-Collared Sparrows (Zonotrichia Capensis) Along an Elevational Gradient. Evolution 63:1593–1605.

Clancy DJ, Hoffmann AA. 1998. Environmental effects on cytoplasmic incompatibility and bacterial load in Wolbachia-infected *Drosophila simulans*. Entomologia Experimentalis et Applicata 86:13–24.

Clayton DA. 2000. Transcription and replication of mitochondrial DNA. Human Reproduction 15 Suppl 2:11–17.

Cogni R, Kuczynski K, Lavington E, Koury S, Behrman EL, O’Brien KR, Schmidt PS, Eanes WF. 2015. Variation in Drosophila melanogaster central metabolic genes appears driven by natural selection both within and between populations. Proceedings of the Royal Society Biological Sciences Series B 282.

Cohen T, Levin L, Mishmar D. 2016. Ancient Out-of-Africa Mitochondrial DNA Variants Associate with Distinct Mitochondrial Gene Expression Patterns. PLOS Genetics 12.

Collinge J, Hoffmann A, McKechnie S. 2006. Altitudinal patterns for latitudinally varying traits and polymorphic markers in *Drosophila melanogaster* from eastern Australia. Journal of Evolutionary Biology 19:473–482.

Consuegra S, John E, Verspoor E, de Leaniz CG. (Consuegra 2015 co–authors). 2015. (Consuegra2015 co-authors). Patterns of natural selection acting on the mitochondrial genome of a locally adapted fish species. Genetics Selection Evolution (Paris) 47:1–10.

Correa CC, Aw WC, Melvin RG, Pichaud N, Ballard JW. 2012. Mitochondrial DNA variants influence mitochondrial bioenergetics in *Drosophila melanogaster*. Mitochondrion 12:459–464.

David JR, Capy P. 1988. Genetic variation of *Drosophila melanogaster* natural populations. Trends in Genetics 4:106–111.

Dobler R, Rogell B, Budar F, Dowling DK. 2014. A meta-analysis of the strength and nature of cytoplasmic genetic effects. Journal of Evolutionary Biology 27:2021–2034.

Doi A, Suzuki H, Matsuura ET. 1999. Genetic analysis of temperature-dependent transmission of mitochondrial DNA in Drosophila. Heredity 82:555–560.

Dowling DK. 2014. Evolutionary perspectives on the links between mitochondrial genotype and disease phenotype. Biochimica et Biophysica Acta 1840:1393–1403.

Dowling DK, Abiega KC, Arnqvist G. 2007. Temperature-specific outcomes of cytoplasmic-nuclear interactions on egg-to-adult development time in seed beetles. Evolution 61:194–201.

Dowling DK, Friberg U, Lindell J. 2008. Evolutionary implications of non neutral mitochondrial genetic variation. Trends in Ecology & Evolution 23:546–554.

Endler JA. 1977. Geographic variation, speciation, and clines. Monographs in Population Biology 10:1–246.

Eyre-Walker A. 2017. Mitochondrial Replacement Therapy: Are Mito-nuclear Interactions Likely To Be a Problem? Genetics 205:1365–1372.

Fontanillas P, Dépraz A, Giorgi MS, Perrin N. (pdf co-authors). 2005. Nonshivering thermogenesis capacity associated to mitochondrial DNA haplotypes and gender in the greater white-toothed shrew, *Crocidura russula*. Molecular Ecology 14:661–670.

Foote AD, Morin PA, Durban JW, Pitman RL, Wade P, Willerslev E, Gilbert MTP, da Fonseca RR. 2011. Positive selection on the killer whale mitogenome. Biology Letters 7:116–118.

Fox J. 2002. An R and S-Plus companion to applied regression. Thousand Oaks, Calif.: Sage Publications.

Frank SA, Hurst LD. 1996. Mitochondria and male disease. Nature 383:224.

Garvin MR, Bielawski JP, Sazanov LA, Gharrett AJ. 2015. Review and meta-analysis of natural selection in mitochondrial complex I in metazoans. Journal of Zoological Systematics and Evolutionary Research 53:1–17.

Gemmell NJ, Metcalf VJ, Allendorf FW. 2004. Mother’s curse: the effect of mtDNA on individual fitness and population viability. Trends in Ecology & Evolution 19:238–244.

Hoekstra LA, Siddiq MA, Montooth KL. 2013. Pleiotropic effects of a mitochondrial-nuclear incompatibility depend upon the accelerating effect of temperature in Drosophila. Genetics 195:1129–1139.

Hoffmann AA, Anderson A, Hallas R. 2002. Opposing clines for high and low temperature resistance in *Drosophila melanogaster*. Ecology Letters 5:614–618.

Hoffmann AA, Weeks AR. 2007. Climatic selection on genes and traits after a 100 year-old invasion: a critical look at the temperate-tropical clines in Drosophila melanogaster from eastern Australia. Genetica 129:133–147.

Hurst LD. 2011. Molecular genetics: The sound of silence. Nature 471:582–583.

James JE, Piganeau G, Eyre-Walker A. 2016. The rate of adaptive evolution in animal mitochondria. Molecular Ecology 25:67–78.

Kearse M, Moir R, Wilson A, Stones-Havas S, Cheung M, Sturrock S, Buxton S, Cooper A, Markowitz S, Duran C, et al. 2012. Geneious Basic: An integrated and extendable desktop software platform for the organization and analysis of sequence data. Bioinformatics 28:1647–1649.

Kimchi-Sarfaty C, Oh JM, Kim IW, Sauna ZE, Calcagno AM, Ambudkar SV, Gottesman MM. 2007. A “silent” polymorphism in the MDR1 gene changes substrate specificity. Science 315:525–528.

Kivisild T, Shen PD, Wall DP, Do B, Sung R, Davis K, Passarino G, Underhill PA, Scharfe C, Torroni A, et al. 2006. The role of selection in the evolution of human mitochondrial genomes. Genetics 172:373–387.

Kofler R, Pandey RV, Schlotterer C. 2011. PoPoolation2: identifying differentiation between populations using sequencing of pooled DNA samples (Pool-Seq). Bioinformatics 27:3435–3436.

Lavington E, Cogni R, Kuczynski C, Koury S, Behrman EL, O’Brien KR, Schmidt PS, Eanes WF. 2014. A Small System-High-Resolution Study of Metabolic Adaptation in the Central Metabolic Pathway to Temperate Climates in *Drosophila melanogaster*. Molecular Biology and Evolution 31:2032–2041.

Levin L, Blumberg A, Barshad G, Mishmar D. 2014. Mito-nuclear co-evolution: the positive and negative sides of functional ancient mutations. Frontiers in Genetics 5:448.

Li H, Stephan W. 2006. Inferring the Demographic History and Rate of Adaptive Substitution in Drosophila. PLOS Genetics 2:e166.

Ma X, Kang J, Chen W, Zhou C, He S. 2015. Biogeographic history and high-elevation adaptations inferred from the mitochondrial genome of Glyptosternoid fishes (Sisoridae, Siluriformes) from the southeastern Tibetan Plateau. BMC Ecology and Evolution 15:233.

Matsuura ET, Tanaka YT, Yamamoto N. 1997. Effects of the nuclear genome on selective transmission of mitochondrial DNA in Drosophila. Genes & Genetic Systems 72:119–123.

Mishmar D, Ruiz-Pesini E, Golik P, Macaulay V, Clark AG, Hosseini S, Brandon M, Easley K, Chen E, Brown MD, et al. 2003. Natural selection shaped regional mtDNA variation in humans. Proceedings of the National Academy of Sciences of the United States of America 100:171–176.

Morales HE, Pavlova A, Joseph L, Sunnucks P. 2015. Positive and purifying selection in mitochondrial genomes of a bird with mitonuclear discordance. Molecular Ecology 24:2820–2837.

Mossman JA, Biancani LM, Zhu CT, Rand DM. 2016. Mitonuclear Epistasis for Development Time and Its Modification by Diet in Drosophila. Genetics 203:463–484.

Musto H, Naya H, Zavala A, Romero H, Alvarez-Valín F, Bernardi G. 2004. Correlations between genomic GC levels and optimal growth temperatures in prokaryotes. FEBS Letters 573:73–77.

Nabholz B, Ellegren H, Wolf JBW. 2013. High Levels of Gene Expression Explain the Strong Evolutionary Constraint of Mitochondrial Protein-Coding Genes. Molecular Biology and Evolution 30:272–284.

Nakamura Y, Gojobori T, Ikemura T. 2000. Codon usage tabulated from international DNA sequence databases: status for the year 2000. Nucleic Acids Research 28:292.

O’Neill SL, Giordano R, Colbert AM, Karr TL, Robertson HM. 1992. 16S rRNA phylogenetic analysis of the bacterial endosymbionts associated with cytoplasmic incompatibility in insects. Proceedings of the National Academy of Sciences of the United States of America 89:2699–2702.

Pfaffl MW, Tichopad A, Prgomet C, Neuvians TP. 2004. Determination of stable housekeeping genes, differentially regulated target genes and sample integrity: BestKeeper--Excel-based tool using pair-wise correlations. Biotechnology Letters 26:509–515.

Porcelli D, Butlin RK, Gaston KJ, Joly D, Snook RR. 2015. The environmental genomics of metazoan thermal adaptation. Heredity 114:502–514.

Quintela M, Johansson MP, Kristjansson BK, Barreiro R, Laurila A. 2014. AFLPs and mitochondrial haplotypes reveal local adaptation to extreme thermal environments in a freshwater gastropod. PLoS One 9:e101821.

Quiros PM, Mottis A, Auwerx J. 2016. Mitonuclear communication in homeostasis and stress. Nature Reviews Molecular Cell Biology 17:213–226.

Rand DM. 2001. The Units of Selection on Mitochondrial DNA. Annual Review of Ecology and Systematics 32:415–448.

Rand DM, Haney RA, Fry AJ. 2004. Cytonuclear coevolution: the genomics of cooperation. Trends in Ecology & Evolution 19:645–653.

Rasmussen R. 2001. Quantification on the LightCycler. In: Meuer S, Wittwer C, Nakagawara K-I, editors. Rapid Cycle Real-Time PCR: Springer Berlin Heidelberg. p. 21–34.

Ruiz-Pesini E, Mishmar D, Brandon M, Procaccio V, Wallace DC. 2004. Effects of purifying and adaptive selection on regional variation in human mtDNA. Science 303:223–226.

Salinas T, Duby F, Larosa V, Coosemans N, Bonnefoy N, Motte P, Marechal-Drouard L, Remacle C. 2012. Co-evolution of mitochondrial tRNA import and codon usage determines translational efficiency in the green alga Chlamydomonas. PLOS Genetics 8:e1002946.

Schattner P, Brooks AN, Lowe TM. 2005. The tRNAscan-SE, snoscan and snoGPS web servers for the detection of tRNAs and snoRNAs. Nucleic Acids Research 33:W686–689.

Sharp PM, Averof M, Lloyd AT, Matassi G, Peden JF. 1995. DNA sequence evolution: the sounds of silence. Philosophical Transactions of the Royal Society of London B Biological Sciences 349:241–247.

She H, Yang QA, Shepherd K, Smith Y, Miller G, Testa C, Mao ZX. 2011. Direct regulation of complex I by mitochondrial MEF2D is disrupted in a mouse model of Parkinson disease and in human patients. Journal of Clinical Investigation 121:930–940.

Silva G, Lima FP, Martel P, Castilho R. 2014. Thermal adaptation and clinal mitochondrial DNA variation of European anchovy. Proceedings of the Royal Society Biological Sciences Series B 281.

Singh RS, Long AD. 1992. Geographic variation in Drosophila: From molecules to morphology and back. Trends in Ecology & Evolution 7:340–345.

Šmarda P, Bureš P, Horová L, Leitch IJ, Mucina L, Pacini E, Tichý L, Grulich V, Rotreklová O. 2014. Ecological and evolutionary significance of genomic GC content diversity in monocots. Proceedings of the National Academy of Sciences of the United States of America 111:E4096–E4102.

Smith C, Heyne S, Richter AS, Will S, Backofen R. 2010. Freiburg RNA Tools: a web server integrating INTARNA, EXPARNA and LOCARNA. Nucleic Acids Research 38:W373–377.

Sorensen JG, Nielsen MM, Loeschcke V. 2007. Gene expression profile analysis of *Drosophila melanogaster* selected for resistance to environmental stressors. Journal of Evolutionary Biology 20:1624–1636.

Sun C, Kong QP, Zhang YP. 2007. The role of climate in human mitochondrial DNA evolution: A reappraisal. Genomics 89:338–342.

Tamura K, Stecher G, Peterson D, Filipski A, Kumar S. 2013. MEGA6: Molecular Evolutionary Genetics Analysis version 6.0. Molecular Biology and Evolution 30:2725–2729.

Toews DP, Brelsford A. 2012. The biogeography of mitochondrial and nuclear discordance in animals. Molecular Ecology 21:3907–3930.

Torres TT, Dolezal M, Schlotterer C, Ottenwalder B. 2009. Expression profiling of Drosophila mitochondrial genes via deep mRNA sequencing. Nucleic Acids Research 37:7509–7518.

Tsujino F, Kosemura A, Inohira K, Hara T, Otsuka YF, Obara MK, Matsuura ET. 2002. Evolution of the A+T-rich region of mitochondrial DNA in the melanogaster species subgroup of Drosophila. Journal of Molecular Evolution 55:573–583.

Wallace DC. 2007. Why Do We Still Have a Maternally Inherited Mitochondrial DNA? Insights from Evolutionary Medicine. Annual Review of Biochemistry 76:781–821.

Weeks AR, McKechnie SW, Hoffmann AA. 2002. Dissecting adaptive clinal variation: markers, inversions and size/stress associations in *Drosophila melanogaster* from a central field population. Ecology Letters 5:756–763.

Williams BR, Van Heerwaarden B, Dowling DK, SgrÒ CM. 2012. A multivariate test of evolutionary constraints for thermal tolerance in *Drosophila melanogaster*. Journal of Evolutionary Biology 25:1415–1426.

Wolff JN, Ladoukakis ED, Enríquez JA, Dowling DK. 2014. Mitonuclear interactions: evolutionary consequences over multiple biological scales. Philosophical Transactions of the Royal Society of London B Biological Sciences 369.

Wolff JN, Tompkins DM, Gemmell NJ, Dowling DK. 2016. Mitonuclear interactions, mtDNA-mediated thermal plasticity, and implications for the Trojan Female Technique for pest control. Scientific Reports 6:30016.

